# GCM: metric-guided clustering by genetic algorithm for correlation-defined modules

**DOI:** 10.64898/2026.07.15.738632

**Authors:** Luis Javier Madrigal-Roca, John K. Kelly

**Affiliations:** Department of Ecology and Evolutionary Biology, University of Kansas

**Keywords:** clustering, genetic algorithm, gene co-expression, gene modules

## Abstract

Gene co-expression analyses identify “good” modules by a correlation criterion. However, standard pipelines detect modules with greedy algorithms that optimize other quantities and only measure correlation afterwards. We present a method called Genetic Clustering by Metric(GCM hereafter), an open-source Python tool that closes this gap by treating module detection as maximum-likelihood inference and solving it globally. In GCM, the correlation objective is, up to a constant and the sample-size factor, the profile log-likelihood of an explicit generative model: a block-diagonal one-factor Gaussian in which each module is a single regulator with equal-magnitude ± loadings. This bases model selection on a principled footing through a genuine BIC/AIC in correlation space. GCM maximizes this likelihood with a memetic genetic algorithm: a population-based search hybridized with a greedy local refinement that reassigns genes after the fact, a move the agglomerative clustering at the core of co-expression pipelines cannot make. Across a replicated noise sweep, GCM reproducibly surpasses hierarchical correlation clustering and k-means with the lowest variance, and an ablation shows the local-search step is responsible; the advantage persists when the number of modules is unknown and when unstructured genes must be ignored. GCM faithfully optimizes geometric indices on the Iris benchmark dataset. For a breast-cancer RNA-seq it recovers coherent modules that predict tumor-versus-normal status. GCM depends only on NumPy and SciPy and exposes one swappable-metric interface with single- and multi-objective modes.

**Author summary:** When biologists group genes by how similarly they are expressed, they usually run a standard clustering method and then score the result with a separate quality measure. The method, however, was never trying to do well on that measure because it optimizes its own internal objective. We built a tool, GCM, that removes this gap: the user picks the quality measure they actually care about, and the tool searches directly for the grouping that scores best on it. The search is performed by a genetic algorithm, a population-based optimizer that mixes and mutates candidate groupings over many generations. GCM includes a purpose-built score for “modules” of co-expressed genes, as well as several widely used geometric scores, and it can balance two competing scores at once to choose how many groups the data support. We show on synthetic data with a known answer, on a textbook dataset, and on real expression data that the tool recovers the intended structure and lets researchers make explicit, and optimize for, their own definition of a good cluster.

## Introduction

High-throughput technologies for profiling gene expression have revolutionized the field of molecular biology, enabling researchers to capture the intricate patterns of gene regulation and unravel the underlying genetic networks. However, the sheer volume and complexity of gene expression data pose significant challenges in extracting meaningful insights [1]. One critical step in this process is the clustering of genes based on their expression profiles, which aims to group together genes exhibiting similar patterns of expression across various experimental conditions or time points [2]. Clustering is a foundational step in computational biology, from grouping co-expressed genes into putative regulatory modules [3, 4] to shedding light on regulatory pathways mechanisms [5], disease-associated genes [6], and potential therapeutic targets [7]. By partitioning a set of objects into groups based on similarity, cluster analysis identifies intrinsic structures in data to gain insight [8, 9].

The dominant practice has two stages. First, a fixed algorithm such as k-means [10] or hierarchical clustering [3] produces a partition. Second,a cluster-validity index like the silhouette [11], the Davies–Bouldin index [12], the Calinski–Harabasz criterion [13], or an information criterion such as BIC [14] is used to *evaluate* that partition, often to choose the number of clusters. The objective the algorithm optimizes (for k-means, within-cluster sum of squares) and the objective by which the result is ultimately judged are therefore distinct. When the two disagree, the reported partition is optimal for neither the user’s stated criterion nor, necessarily, the underlying biology.

This gap is particularly consequential in gene co-expression analysis. There, the notion of a good module is intrinsically correlational: genes in a module should be strongly co-expressed across conditions, regardless of their absolute expression levels or Euclidean geometry. Tools such as WGCNA [15], Clust [16], and UNCLES [17] encode this intuition through correlation networks or consensus procedures, and comparative studies show that module-detection performance depends strongly on how the objective is framed [18]. Yet the correlation-based quality of the final modules is still, in most pipelines, assessed only after the clustering algorithm has finished optimizing a different, geometric objective.

We argue for inverting the workflow by making the validity metric itself the optimization target. Concretely, the user specifies the scalar criterion that defines a good partition for their problem, and a search procedure evolves a partition to optimize that criterion directly. This is attractive whenever the criterion of interest is (i) not what standard algorithms optimize, (ii) non-differentiable or combinatorial, or (iii) most naturally expressed as a trade-off between competing concerns—fit versus parsimony, compactness versus separation. Direct optimization of clustering criteria has a long history in evolutionary computation [19, 20], but it is rarely packaged as a practical, dependency-light tool aimed at the co-expression use case and built around a correlation objective.

While traditional numerical optimizers—such as Newton-Raphson, Powell’s method, or the Simplex algorithm—are highly effective for continuous parameter estimation, they are fundamentally ill-suited for this clustering framework. These methods require a continuous search space and differentiable objective functions to calculate valid step directions. In contrast, identifying optimal partitions is an inherently discrete, combinatorial problem. Our objective functions are highly discontinuous with respect to categorical cluster labels, resulting in a rugged fitness landscape riddled with local optima where gradient-based methods become trapped. Furthermore, while stochastic Bayesian approaches (such as Reversible Jump MCMC) could theoretically navigate this discrete landscape by jumping between partition states, the computational time cost of sufficiently sampling such a vast combinatorial space is prohibitive for large datasets. The Genetic Algorithm provides the necessary robustness to directly evolve discrete partition vectors, efficiently navigating the non-differentiable landscape via crossover and mutation without the extreme computational overhead of exhaustive Bayesian sampling or the restrictive assumptions of gradient-based methods.

Here we present GCM (Genetic Clustering by Metric), an open-source Python package that clusters data by directly optimizing a chosen cluster-validity metric with a memetic genetic algorithm (GA), which is an evolutionary algorithm hybridized with a problem-specific local search that refines each candidate solution within its own “lifetime,”. This strategy was introduced by Moscato [21] and surveyed by Neri and Cotta [22]. Genetic algorithms (GAs) could be a promising solution for clustering tasks [23], particularly in the context of gene expression data [9]. Other attempts have been made to use evolutionary algorithms in this or similar contexts, with overall promising results [24–27]. GAs are optimization algorithms that mimic the principles of natural selection and evolution, making them well-suited for complex, multi-objective problems where traditional methods may struggle to find optimal solutions [28].

GCM makes three contributions. First, we show that the project’s correlation objective is, up to a constant and the sample-size factor, the profile log-likelihood of an explicit block-diagonal one-factor Gaussian, in which each module is a single regulator with equal-magnitude *±* loadings, so that module detection becomes maximum-likelihood inference and the number of modules is chosen by a genuine BIC/AIC in correlation space, not a heuristic. Second, we hybridize the GA with an efficient memetic local search and show, with a replicated benchmark, that this global optimizer reproducibly surpasses the greedy agglomerative clustering at the core of standard co-expression pipelines, with an ablation attributing the gain to the local-search step. Third, the implementation is deliberately minimal—depending only on NumPy [29] and SciPy [30], with internal k-means++ [31] and correlation-distance hierarchical seeders, behind one swappable-metric interface that also exposes standard geometric indices and a multi-objective (NSGA-II [32]) mode. We evaluate GCM on synthetic data across a noise sweep with replication, on the classic Iris benchmark [33], and on a breast-cancer RNA-seq dataset.

## Materials and methods

### Overview and problem statement

GCM clusters *n elements* (the rows of a data matrix **X** *∈* ***ℝ***^*n×d*^; in co-expression analysis, *n* genes measured across *d* samples) by searching the space of partitions for one that optimizes a user-chosen validity metric. A partition is represented as an integer label vector ***ℓ*** *∈ {*0, 1, …, *K}*^*n*^, where *ℓ*_*i*_ = *c* assigns element *i* to cluster *c* and the optional label 0 marks an element as unassigned. The user fixes an upper bound *g*_max_ on the number of clusters. Each candidate partition ***ℓ*** is scored by one or more metric functions *m*(***ℓ***) *∈* ***ℝ***; the GA evolves a population of partitions to optimize these scores in their natural direction (maximize or minimize).

### Performance-target metrics

GCM treats every metric as an interchangeable “performance target” or fitness measure [28]. Table 1 lists the built-in targets, the input format each one is design to work with, and its optimization direction. They fall into two families: correlation-based objectives designed for modular data, and standard geometric/probabilistic indices.

**Table 1.**
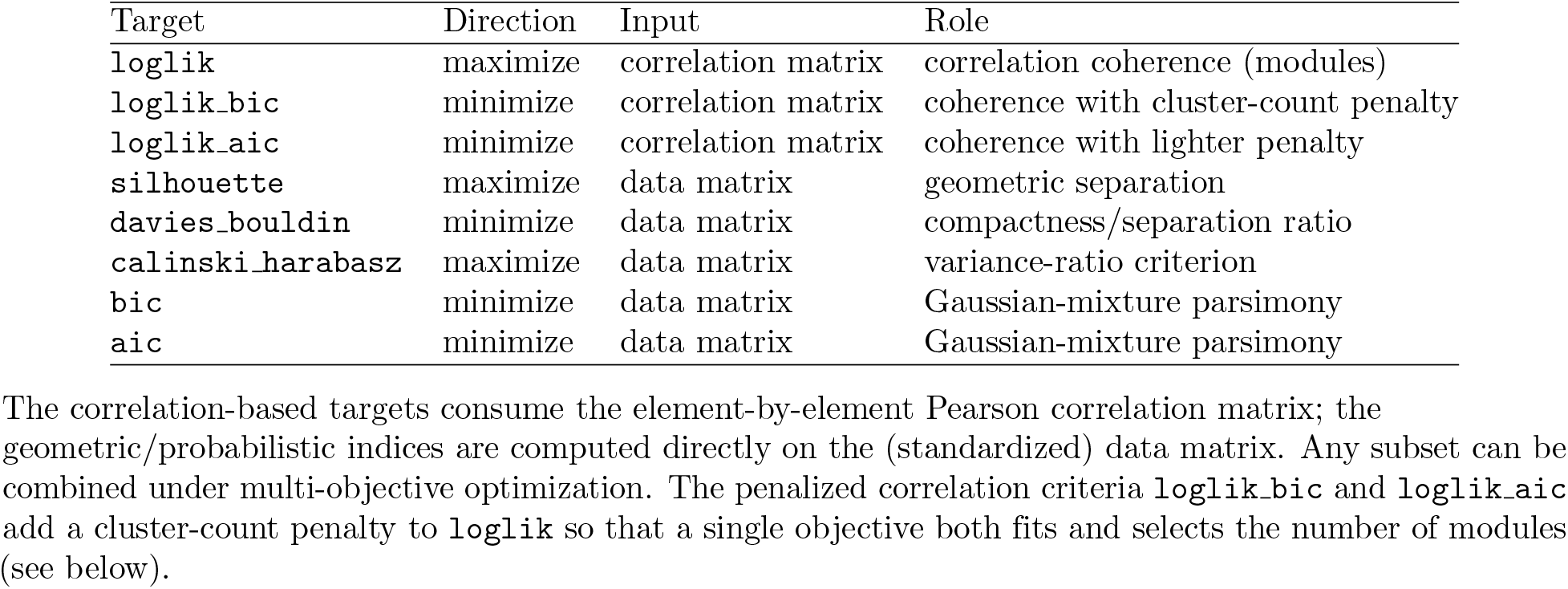
The built-in performance-target metrics.

#### A generative model for correlation modules

The correlation objective is the maximum log-likelihood of an explicit generative model, which we state here and derive in full in S1 Appendix. Let the data matrix **X** ∈ ℝ^*n×d*^ hold *n* genes measured over *d* samples, row-standardized. We model each *sample* (column) as an independent draw from a multivariate Gaussian over genes, *N* (**0**, Σ), whose correlation matrix Σ is *block-diagonal* by the partition ***ℓ***: genes in the same module are correlated, genes in different modules independent. Within a module *s* we adopt the canonical single-regulator (one-factor) structure with equal-magnitude loadings of arbitrary sign, *x*_*g*_ = *σ*_*g*_*f*_*s*_ + *ε*_*g*_ with *σ*_*g*_ *∈ {±λ*_*s*_*}*, so that any two genes in the module share |corr| = *ρ*_*s*_. This is exactly the core assumption of a co-expression module (one regulator, up- or down-regulating its targets).

#### The likelihood reduces to the project’s objective

Let **R** be the Pearson correlation matrix between genes and, for module *s* of size *n*_*s*_, let

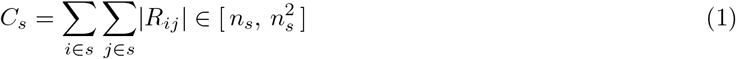

(the unit diagonal is included). As shown in S1 Appendix, the maximum-likelihood common coherence of a module is its mean absolute off-diagonal correlation, 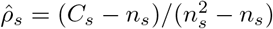, and at that estimate the trace term of the Gaussian log-likelihood collapses to the module dimension. The profile log-likelihood of a partition is therefore, up to a partition-independent constant,

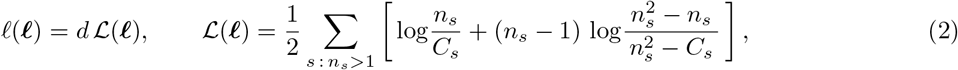

where *L* is precisely 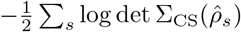, the equicorrelation log-determinant. Maximizing *L* thus maximizes the model likelihood. Each summand is monotone in module coherence: as 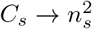 the cluster is perfectly correlated (best fit) and as *C*_*s*_ → *n*_*s*_ it is incoherent (contributing zero); singletons contribute nothing. A perfectly correlated cluster is awarded a large finite reward rather than +*∞* for numerical stability.

#### Model selection: information criteria

The raw likelihood favours more modules, since splitting a coherent module into coherent sub-modules does not necessarily lower *L*. Because each module carries a single coherence parameter *ρ*_*s*_, the model with *K* modules has *K* free parameters, and the standard information criteria over the *d* samples are

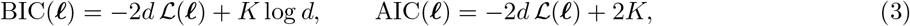

to be minimized (targets `loglik_bic`, `loglik_aic`). The likelihood scales with the sample size *d* while the penalty does not, so more samples correctly license more modules. This is distinct from the geometric bic/aic targets, whose Euclidean-Gaussian assumption conflicts with the absolute-correlation objective when modules contain anti-correlated members, which is precisely the regime in which the correlation criteria are the right choice.

#### Geometric and probabilistic indices

For data whose clusters are defined geometrically, GCM provides four standard indices computed on **X**: the mean silhouette coefficient [11] (maximize; range [−1, 1]), the Davies–Bouldin index [12] (minimize), the Calinski–Harabasz variance-ratio criterion [13] (maximize), and the Bayesian and Akaike information criteria [14, 34] under a spherical-Gaussian-per-cluster model with pooled variance (both minimize). The BIC/AIC log-likelihood follows the x-means formulation [35]; for a partition with *k* clusters in *d* dimensions the model has *p* = *kd* + (*k*− 1) + 1 free parameters (centroids, mixture weights, and one pooled variance), and 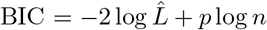, AIC 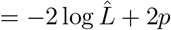. Unlike the correlation objective, BIC and AIC reward parsimony explicitly and so can be used for model selection on geometrically compact data.

The correlation-based targets consume the element-by-element Pearson correlation matrix; the geometric/probabilistic indices are computed directly on the (standardized) data matrix. Any subset can be combined under multi-objective optimization. The penalized correlation criteria `loglik_bic` and `loglik_aic` add a cluster-count penalty to `loglik` so that a single objective both fits and selects the number of modules (see below).

### Data preparation

Input is a numeric matrix (CSV/TSV or array) with rows as elements and columns as features; header rows and index columns are auto-detected. By default the data are standardized to zero mean and unit variance. For the correlation objective, standardization is applied per row (per element/gene), which is the regime under which the Pearson correlation between rows is meaningful; for the geometric indices, per-feature standardization is the appropriate choice. The element-by-element correlation matrix **R** is computed once and reused across all evaluations of *L*.

### Dimensionality-reduction visualization

Visual inspection of clustering results is essential for communicating and sanity-checking a partition. GCM provides built-in tools for projecting the clustered data into two (or three) dimensions, with the cluster labels shown as colours. Two embedding methods are always available (no extra dependencies): *principal component analysis* (PCA), a linear projection preserving maximal variance, and *correlation multidimensional scaling* (correlation MDS), a classical MDS embedding [36] on the correlation distance matrix 1 − |**R**|. The second method is particularly informative for the correlation objective because it projects the elements into the same distance space the log-likelihood operates in: if modules are well-separated in correlation, they appear well-separated in the MDS embedding. For users with additional packages installed, t-SNE [37] and UMAP [38] are also supported and may better resolve local neighbourhood structure in large or high-dimensional datasets. These tools are exposed both through the Python API (`reduce_dimensions` and `plot_clusters`) and the command line (`gcmrk visualize`).

### Genetic algorithm

GCM evolves a population of partitions with a self-contained GA (Fig 3). The components are:

1. **Initialization**. The initial population is seeded with three kinds of partition across candidate cluster counts *k* = 2, …, *g*_max_: k-means++ partitions (a small internal NumPy k-means with multiple restarts); *correlation-aware* partitions obtained by agglomerative (average- and complete-linkage) hierarchical clustering of the correlation distance 1 − |**R**|, the classical co-expression clustering strategy; and random partitions for diversity. User-supplied seeds (e.g. the output of WGCNA or Clust) can also be injected. The correlation-aware seeds are important for the loglik objective: Euclidean k-means splits strongly anti-correlated elements that are coherent under the absolute-correlation criterion, whereas the correlation-distance seeds already respect the structure the GA is asked to optimize.
2. **Evaluation**. Each partition is scored on the chosen target(s); a cache keyed on the label vector avoids recomputing identical partitions.
3. **Selection and variation**. In *weighted* mode the targets are combined into a single fitness through direction-aware weights after a bounded squashing transform that places metrics of different scales on a comparable footing, and elitist tournament selection is applied. In *nsga2* mode the targets are treated as competing objectives and NSGA-II [32] fast non-dominated sorting with crowding distance is used. Variation is two-point crossover on label vectors plus a reassignment mutation that moves a small fraction of elements to random clusters.
4. **Repair**. After every operator a partition is relabelled to consecutive integers and constrained to have between 2 and *g*_max_ clusters, so that all validity metrics remain well defined (degenerate single-cluster or empty-cluster partitions are repaired).
5. **Memetic local search**. For the correlation objective, the seed partitions and the elite of each generation are refined by a greedy hill-climb that repeatedly applies the single element reassignment most improving the objective until none remains (a memetic, Lamarckian GA). The gain of a move touches only the source and target modules, so it is computed incrementally in *O*(module size) from *C*_*s*_ (Eq (1)); a full sweep is *O*(*n*^2^). This step is decisive: it performs exactly the move that agglomerative clustering—used here only for seeding—cannot, namely reassigning an element after an early merge, and as shown in the Results it is what lets the optimizer surpass hierarchical clustering. When the objective is a penalized criterion (Eq (3)) the hill-climb optimizes that criterion, so it also merges spurious modules.

For multi-objective runs GCM returns the full Pareto front and, as a single representative partition, the knee point that maximizes the sum of min–max normalized objectives across the front. Unless stated otherwise, all experiments use a population of 200 evolved for 120 generations, two k-means restarts per candidate *k*, crossover probability 0.8, mutation probability 0.2 with per-gene reassignment probability 0.05, tournament size 3, and elitism 5, with a fixed random seed for reproducibility.

### Choosing the number of clusters

Because *L* (Eq (2)) has no parsimony term (penalization against more clusters), GCM offers three routes to the right number of clusters. (i) *Constrain g*_max_ to the expected number of modules. (ii) *Optimize a penalized correlation criterion* (`loglik_aic` or `loglik_bic`, Eq (3)) as a single objective under a loose *g*_max_; this selects *K* within correlation space and is our recommended default when *K* is unknown. Equivalently, one can scan *g*_max_ with loglik and pick the *K* that minimizes Eq (3). (iii) *Explore an explicit trade-off* by optimizing loglik jointly with a parsimony target under NSGA-II and reading the knee of the Pareto front; pairing with the geometric bic is informative only when the modules are also geometrically compact, whereas the penalized correlation criteria avoid that assumption.

### Benchmark datasets and evaluation

We evaluate on three datasets with known ground truth and one empirical expression dataset.

*Synthetic-Easy* and *Synthetic-Hard* are generated by GCM’s modular simulator: each module is driven by a shared latent factor (with random *±* 1 loadings, so within-module genes are strongly correlated in absolute value while genes in different modules are not), with per-entry Gaussian noise added and rows finally standardized. Synthetic-Easy has three equal modules (75 genes across 80 samples) at low noise, giving strong separation (mean within-module |*r*| = 0.66 versus between-module 0.07). Synthetic-Hard has five unequal modules (15, 20, 25, 30, 10 genes; 100 genes across 50 samples) at higher noise, giving weak separation (within 0.33 versus between 0.12).

*Iris* [33] is the classic 150-sample, 4-feature, 3-class benchmark; it is low-dimensional and geometric rather than correlational, and we use it to show that the same engine, driven by geometric targets, faithfully optimizes those indices on data outside the co-expression regime.

*Empirical data*. We apply GCM to GSE183947 [39], a breast cancer RNA-seq dataset of 60 samples (30 tumours and 30 patient-matched adjacent-normal tissues) with 20246 genes quantified as FPKM. After log_2_ transformation we cluster the 200 most variable genes into co-expression modules, and assess each module both by its internal correlation coherence and by whether its eigengene (the leading principal component of the module’s expression across samples) distinguishes tumour from normal tissue (Welch *t*-test), linking the recovered modules to the clinical phenotype.

*Decisive benchmark*. To test whether globally optimizing the block likelihood actually improves on greedy alternatives, we run a replicated benchmark on the five-module design over a noise sweep *σ ∈ {*0.7, 1.0, 1.5, 2.0 *}*, with 30 random datasets per noise level. We compare: GCM (memetic); GCM with the local-search step disabled (an ablation isolating its contribution); agglomerative clustering on 1 − |**R**| with average and with complete linkage (the algorithmic core of correlation-network methods such as WGCNA), cut at the true *k*; and k-means. We also evaluate model selection with *k* unknown (GCM/loglik aic versus hierarchical clustering whose cut minimizes the same penalized criterion) and robustness to unstructured “noise” genes that belong to no module.

We use the adjusted Rand index (ARI) [40] wirsanskyHGA2020; for the single-dataset summaries (Table 3) we also report Hungarian-matched accuracy. Results are reported as mean *±* SD over the random datasets. All code to reproduce the experiments, figures, and tables is provided with the package.

**Table 2.** Module recovery (ARI, mean *±* SD over 30 datasets) on the five-module design across a noise sweep, with the number of clusters fixed to the truth.

| Method | $\sigma = 0.7$ | $\sigma = 1.0$ | $\sigma = 1.5$ | $\sigma = 2.0$ |
| --- | --- | --- | --- | --- |
| GCM (memetic) | 1.00±0.00 | 1.00±0.00 | 0.94±0.05 | 0.50±0.17 |
| GCM (no local search) | 1.00±0.00 | 1.00±0.01 | 0.88±0.12 | 0.53±0.15 |
| hierarchical (average) | 1.00±0.00 | 1.00±0.01 | 0.80±0.14 | 0.44±0.12 |
| hierarchical (complete) | 1.00±0.00 | 0.98±0.06 | 0.55±0.16 | 0.16±0.10 |
| k-means | 0.37±0.03 | 0.36±0.03 | 0.31±0.05 | 0.18±0.06 |

**Table 3.** Benchmark results: clusters recovered (*k*), adjusted Rand index (ARI), and Hungarian-matched accuracy against ground truth.

| Dataset | Method | Target | Setting | $k$ | ARI | Acc. |
| --- | --- | --- | --- | --- | --- | --- |
| Synthetic-Easy | k-means | WCSS | k=3 | 3 | 0.220 | 0.587 |
|  | GCM | loglik | known k=3 | 3 | 1.000 | 1.000 |
|  | GCM | loglik | loose g_max=10 | 3 | 1.000 | 1.000 |
|  | GCM | loglik_bic | model sel. g_max=10 | 3 | 1.000 | 1.000 |
|  | GCM | loglik_aic | model sel. g_max=10 | 3 | 1.000 | 1.000 |
| Synthetic-Hard | k-means | WCSS | k=5 | 5 | 0.269 | 0.430 |
|  | GCM | loglik | known k=5 | 5 | 0.986 | 0.990 |
|  | GCM | loglik | loose g_max=10 | 6 | 0.903 | 0.900 |
|  | GCM | loglik_bic | model sel. g_max=10 | 6 | 0.903 | 0.900 |
|  | GCM | loglik_aic | model sel. g_max=10 | 6 | 0.903 | 0.900 |
|  | GCM/nsga2 | loglik+bic | nsga2 g_max=10 | 5 | 0.332 | 0.550 |
| Iris | k-means | WCSS | k=3 | 3 | 0.592 | 0.813 |
|  | GCM | silhouette | g_max=6 | 2 | 0.568 | 0.667 |
|  | GCM | calinski_harabasz | g_max=6 | 2 | 0.568 | 0.667 |
|  | GCM/nsga2 | silhouette+davies_bouldin | nsga2 g_max=8 | 3 | 0.558 | 0.660 |
| GSE183947 | GCM | loglik_aic | top200, g_max=12 | 12 | – | – |

## Results

### Recovering correlated modules on synthetic data

On Synthetic-Easy, GCM driven by the correlation log-likelihood (with *g*_max_ set to the true count) recovers the planted three-module structure perfectly (ARI = 1.00), whereas the k-means baseline—given the same true *k*—fails badly (ARI = 0.22). The gap is diagnostic: because module loadings carry random signs, each module contains strongly anti-correlated genes that are coherent under the absolute-correlation objective but lie far apart in Euclidean space, so k-means splits them. The GA converges within the first few dozen generations (Fig 3). On Synthetic-Hard, where weak separation and higher noise make the problem substantially harder, GCM still recovers the five modules well (ARI = 0.99) while k-means remains poor (ARI = 0.27). The correlation-aware hierarchical seeding (Methods) is important here: it supplies the GA with starting partitions that already respect the absolute-correlation structure, which a Euclidean seeder cannot.

### Global optimization beats greedy clustering, and local search is why

The central methodological question is whether globally optimizing the block likelihood improves on the greedy agglomerative clustering it is seeded from. The replicated noise sweep (Fig 1, Table 2) shows that the answer is yes, decisively and reproducibly. At the hard noise level (*σ* = 1.5, mean over 30 datasets), GCM attains ARI 0.94 versus 0.80 for average-linkage hierarchical clustering at the true *k* and 0.31 for k-means, and its variance across datasets is the smallest of all methods, so the advantage is consistent rather than occasional. The gap widens as noise grows. An ablation pins the cause on the memetic local search: disabling it drops GCM to ARI 0.88 (Fig 2). The genetic operators and correlation-aware seeds alone already match or modestly exceed hierarchical clustering, but it is the hill-climb with the designed reassignment of elements *a posteriori* (a move greedy agglomeration cannot make) the feature that produces the decisive margin and the low variance.

**Fig 1.**
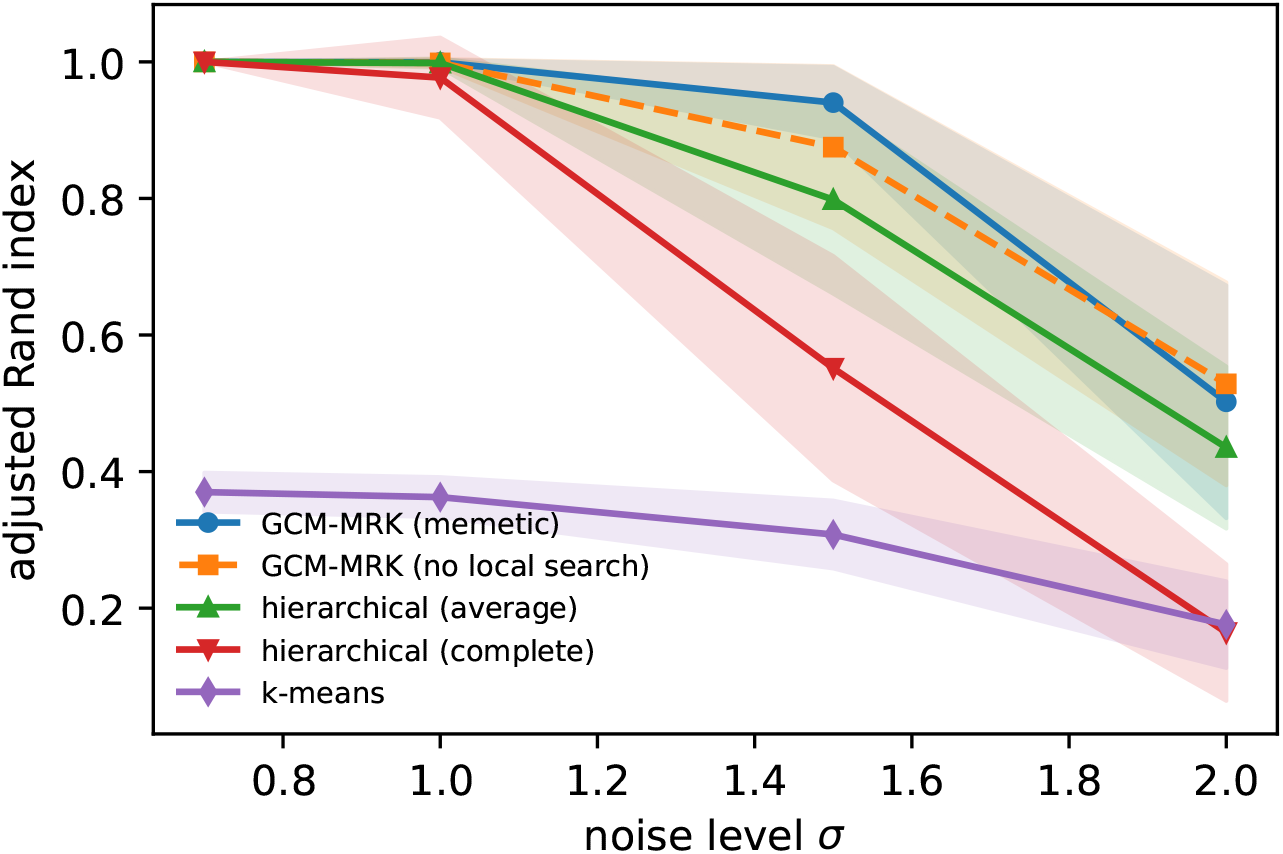
Module recovery versus noise. Adjusted Rand index (mean, bands *±* SD over 30 datasets) at the true number of clusters. GCM (memetic) dominates hierarchical clustering and k-means across the noise range and is the most consistent; the ablation without local search lies in between.

**Fig 2.**
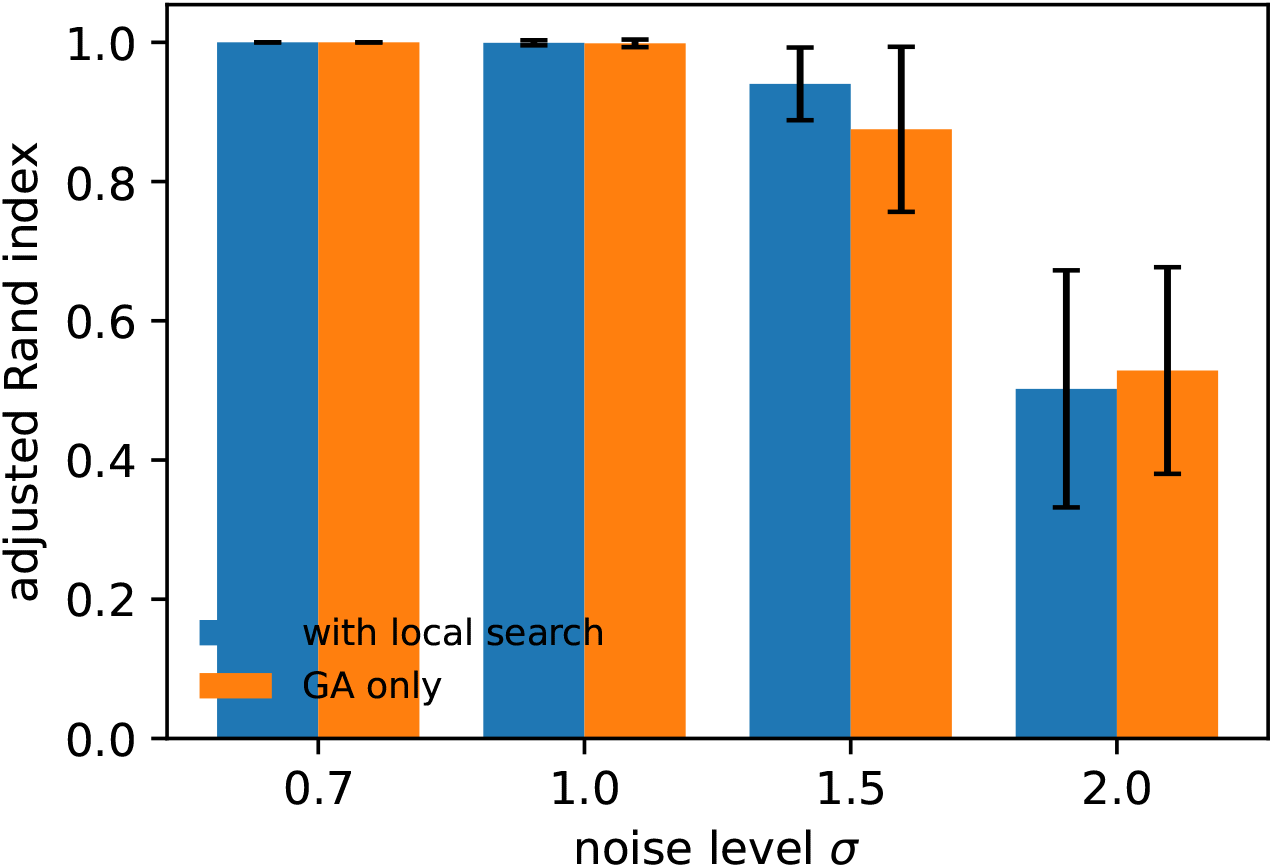
Local search is the decisive ingredient. Recovery with and without the memetic hill-climb across the noise sweep; the refinement that reassigns elements after seeding accounts for most of GCM’s advantage over greedy clustering.

**Fig 3.**
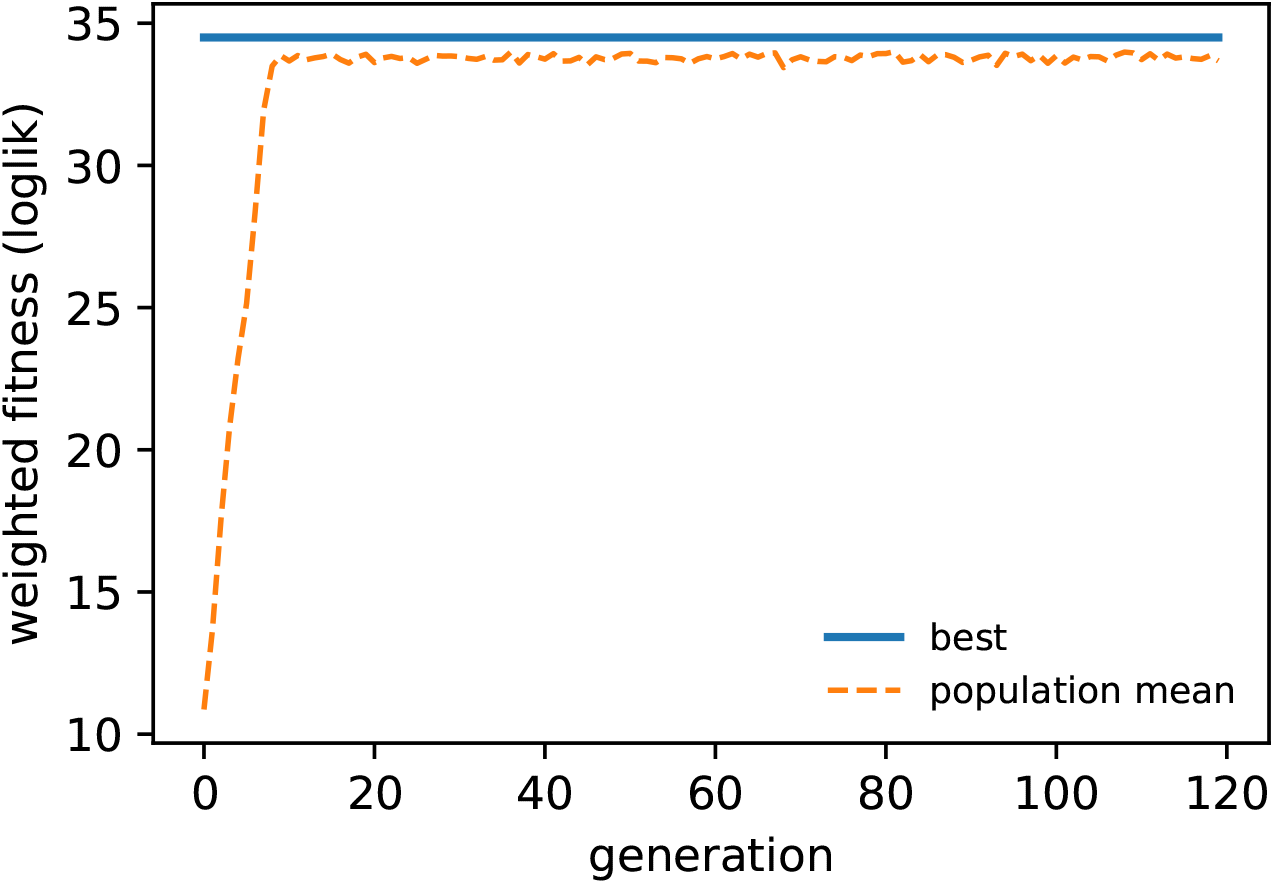
Convergence of the genetic algorithm on Synthetic-Easy. Best and population-mean weighted fitness (correlation log-likelihood) across generations. The optimizer reaches a high-quality partition within the first few dozen generations and the elite fitness is monotone non-decreasing by construction.

### Selecting the number of clusters without knowing it

When only loglik is optimized under a loose *g*_max_, the correlation objective can over-segment, since Eq (2) carries no parsimony term. The effect, however, is mild once the likelihood is optimized well: on Synthetic-Easy the optimum already coincides with the true partition (*k* = 3), and on Synthetic-Hard a loose bound yields only *k* = 6 (true 5; Table 3), one module above the truth. The penalized correlation criteria (Eq (3)) keep selection near the truth within a single objective: on Synthetic-Easy `loglik_aic` returns *k* = 3 at ARI = 1.00, and on this Synthetic-Hard dataset *k* = 6 at ARI = 0.90. The decisive test is replication: over 30 datasets `loglik_aic` selects 6.7 modules on average at ARI 0.87, closer to the truth and more accurate than hierarchical clustering whose cut minimizes the same criterion (7.5 modules, ARI 0.83; Table 4). So, the advantage of global optimization persists when the module count must be inferred. The fit–parsimony trade-off is also visible by scanning *g*_max_ (Fig 4): `loglik` rises and plateaus with *k* while the penalized criteria turn over near the true number of modules.

**Table 4.**
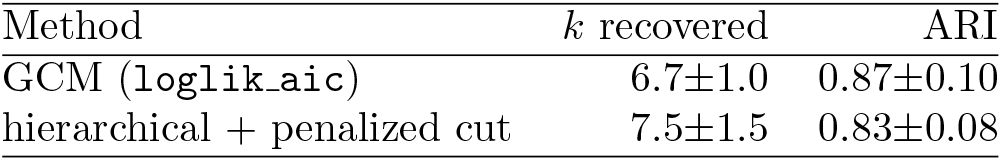
Model selection with *k* unknown. (mean *±* SD over 30 datasets, Synthetic-Hard design, true *k* = 5). Both methods use the same penalized correlation criterion; they differ only in optimizer.

**Fig 4.**
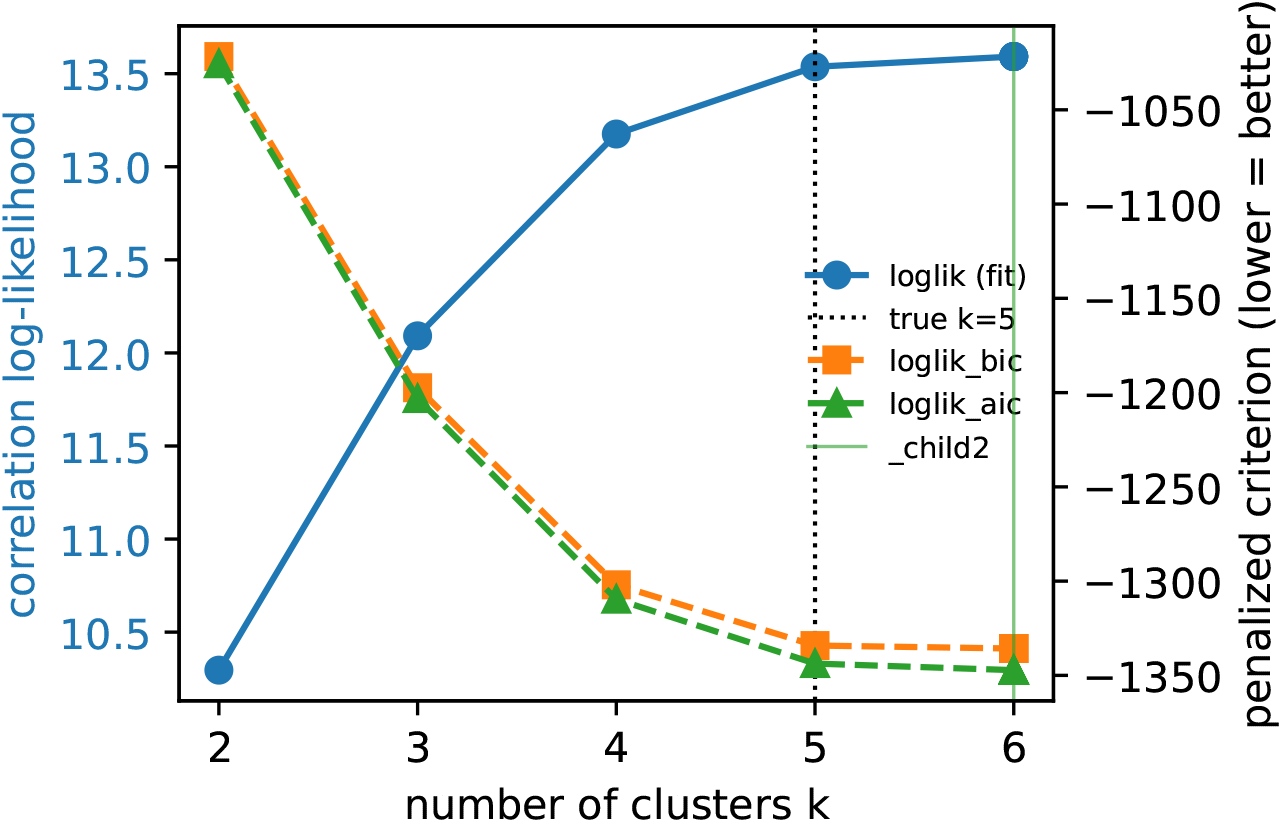
Model selection on Synthetic-Hard. The raw correlation log-likelihood (left axis) increases monotonically as clusters are added—the over-segmentation tendency—while the penalized criteria (right axis, lower is better) turn over near the true number of modules. The `loglik_aic` minimum identifies a parsimonious, well-fitting partition without the number of clusters being specified in advance.

For users who prefer to inspect an explicit trade-off rather than commit to a single penalized criterion, the multi-objective mode optimizes `loglik` against a parsimony target under NSGA-II and returns the full Pareto front (Fig 5). We note that pairing `loglik` with the *geometric* BIC is less effective on these data, because BIC’s Euclidean-Gaussian assumption conflicts with the absolute-correlation objective when modules contain anti-correlated members—which is exactly why the penalized *correlation* criteria are preferable for module selection.

**Fig 5.**
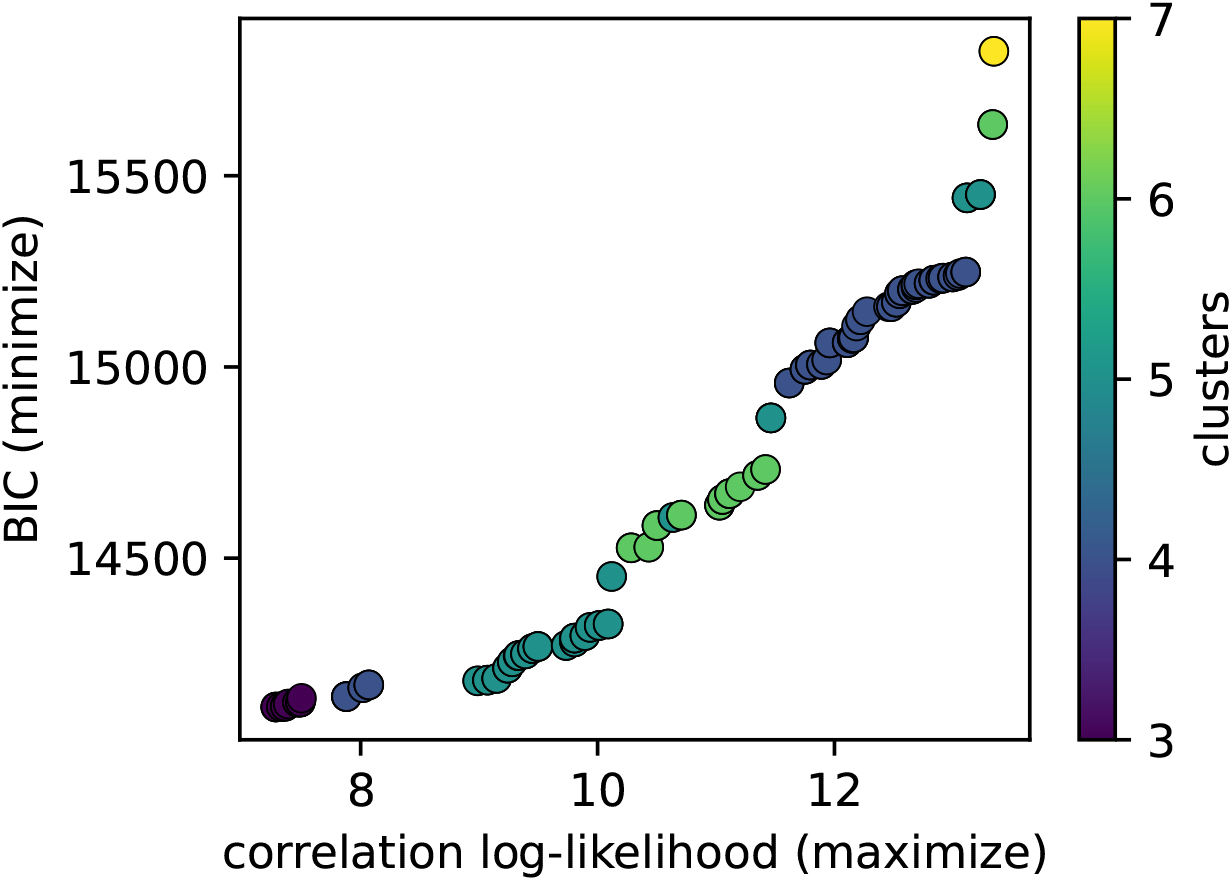
Fit–parsimony Pareto front on Synthetic-Hard. Each point is a non-dominated partition from a single NSGA-II run optimizing the correlation log-likelihood (maximized) against the geometric BIC (minimized); colour encodes the number of clusters. The multi-objective mode exposes the trade-off explicitly for interactive exploration.

True *k* is 3 (Synthetic-Easy, Iris) and 5 (Synthetic-Hard); GSE183947 has no ground-truth labelling (ARI/accuracy not applicable). “known *k*” fixes *g*_max_ to the true value; “loose *g*_max_” and “model sel.” use a loose bound, the latter with a penalized correlation criterion. k-means is given the true *k* and 10 restarts. GA settings as in Methods; fixed random seed.

### Robustness to unstructured genes

Real gene sets contain many genes that belong to no module. When the five-module design is embedded among unstructured “noise” genes and the methods must both find the modules and not be derailed by the noise, GCM retains ARI 0.96 against 0.53 for hierarchical clustering (30 datasets). The likelihood isolates the incoherent genes rather than letting them pollute real modules, whereas greedy linkage forces them into the nearest module and degrades the recovered structure.

### Optimizer validation on geometric data: Iris

We include Iris not as a recovery benchmark—it is geometric rather than correlational, outside the correlation objective’s domain—but to verify that the GA is a faithful, competent optimizer of *any* geometric index it is given. It is: driven by silhouette it returns a partition with silhouette 0.58, matching exactly the optimum found by k-means at *k* = 2 (0.58), so the optimizer reaches the index’s best value rather than a local one.

That this partition has *k* = 2 rather than three clusters is a property of the *index*, not the optimizer: on Iris the three-species labelling has silhouette only 0.38, *below* both the two-cluster split (0.58) and the best three-cluster k-means partition (0.46), because *versicolor* and *virginica* overlap so heavily that merging them is geometrically cleaner (Fig 6). No silhouette-maximizing method can return the three species here; GCM correctly returns what the criterion prefers (ARI = 0.57 against the species, matching k-means at the same *k*). The lesson is that the choice of criterion, not the optimizer, determines the answer, which is precisely the control GCM gives the user, and the reason the correlation objective matters for data where a geometric index would be the wrong question.

**Fig 6.**
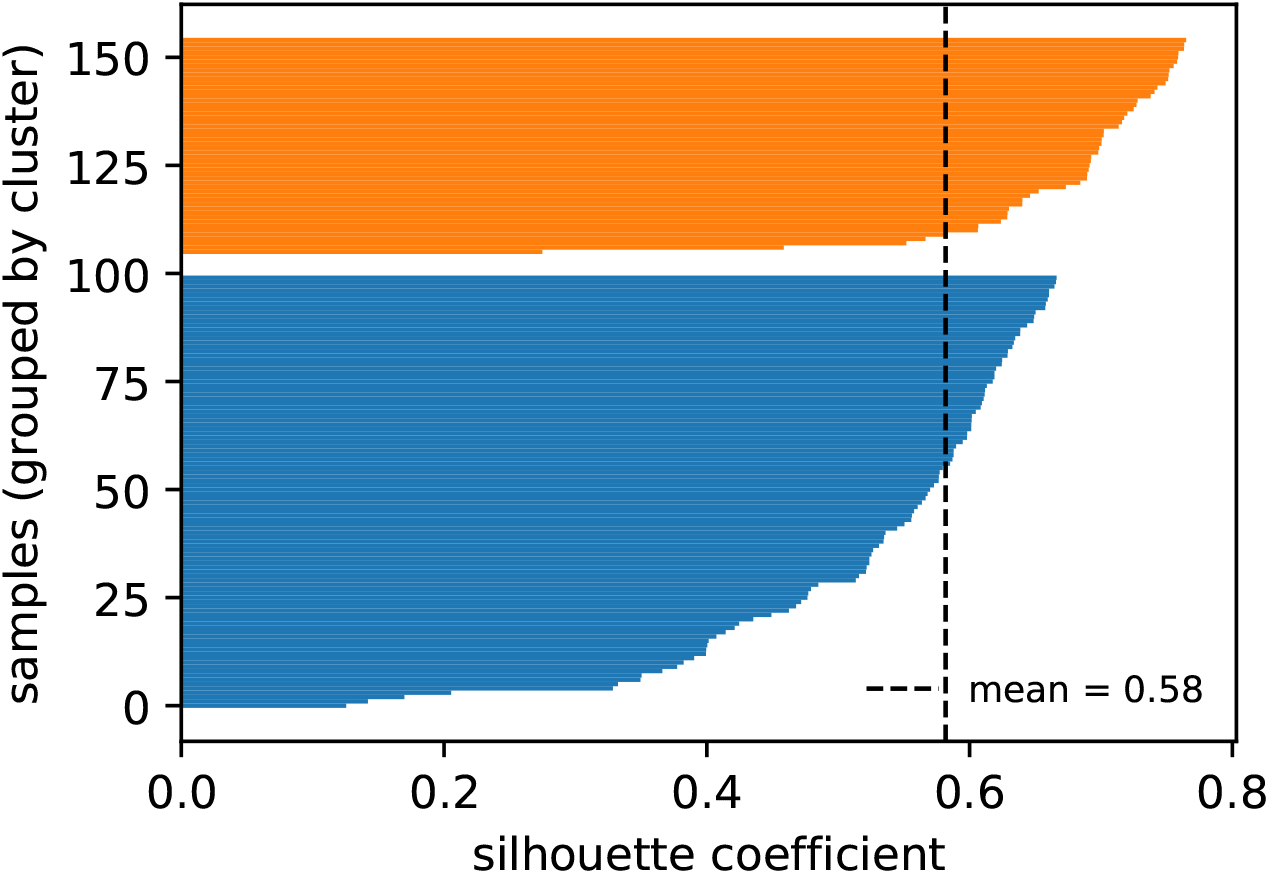
Silhouette profile of the Iris clustering recovered by GCM. Per-sample silhouette coefficients grouped by recovered cluster; the dashed line marks the mean. One cluster is cleanly separated (uniformly high silhouette) while the other two overlap, the expected structure for the Iris species.

### Empirical co-expression modules: breast cancer

Applied to the 200 most variable genes of GSE183947 with the penalized correlation criterion loglik aic, GCM resolves 12 co-expression modules. These are strongly coherent: the mean within-module absolute correlation is 0.59, against a background of 0.31 over all gene pairs. Crucially, the modules are biologically meaningful with respect to the tumour/normal axis that the clustering never saw: 7 of the 12 module eigengenes differ significantly between tumour and patient-matched normal tissue (Welch *t*-test, Benjamini–Hochberg *q <* 0.05; Fig 7, Table 5). Because the genes were selected only for variability and clustered only by mutual correlation, the recovery of phenotype-associated modules indicates that the correlation objective captures co-regulation structure that tracks the underlying biology.

**Table 5.**
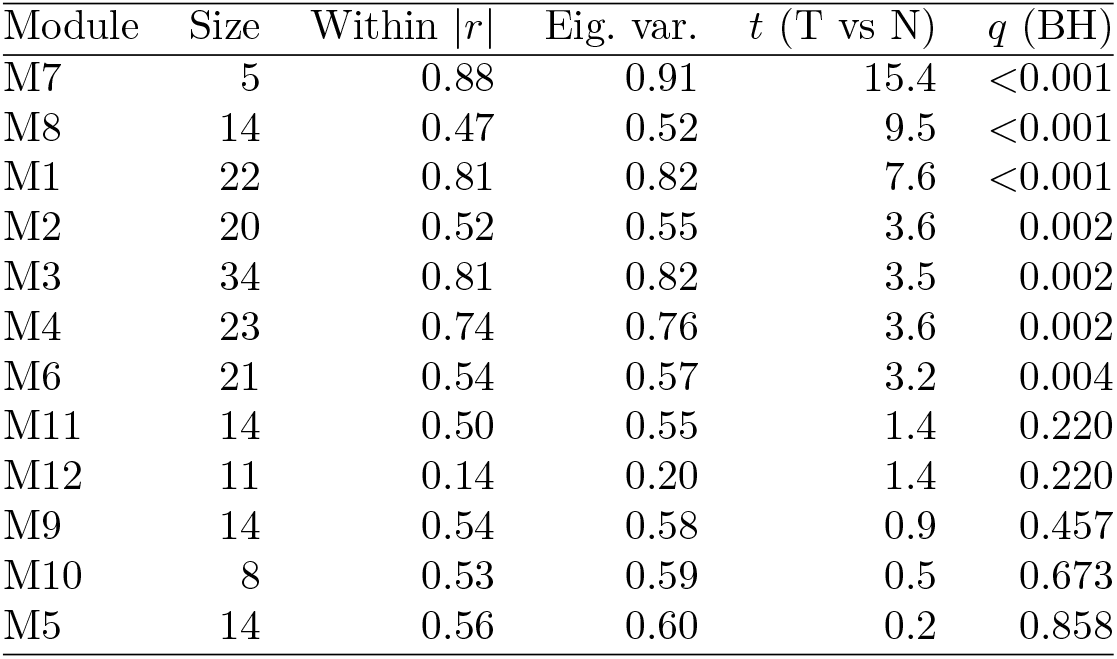
Co-expression modules recovered from GSE183947. Size, mean within-module absolute correlation, variance explained by the module eigengene, and the tumour-vs-normal eigengene contrast (*t* statistic and Benjamini–Hochberg-adjusted Welch *q*-value).

| Module | Size | Within $\vert r\vert $ | Eig. var. | $t$ (T vs N) | $q$ (BH) |
| --- | --- | --- | --- | --- | --- |
| M7 | 5 | 0.88 | 0.91 | 15.4 | <0.001 |
| M8 | 14 | 0.47 | 0.52 | 9.5 | <0.001 |
| M1 | 22 | 0.81 | 0.82 | 7.6 | <0.001 |
| M2 | 20 | 0.52 | 0.55 | 3.6 | 0.002 |
| M3 | 34 | 0.81 | 0.82 | 3.5 | 0.002 |
| M4 | 23 | 0.74 | 0.76 | 3.6 | 0.002 |
| M6 | 21 | 0.54 | 0.57 | 3.2 | 0.004 |
| M11 | 14 | 0.50 | 0.55 | 1.4 | 0.220 |
| M12 | 11 | 0.14 | 0.20 | 1.4 | 0.220 |
| M9 | 14 | 0.54 | 0.58 | 0.9 | 0.457 |
| M10 | 8 | 0.53 | 0.59 | 0.5 | 0.673 |
| M5 | 14 | 0.56 | 0.60 | 0.2 | 0.858 |

**Fig 7.**
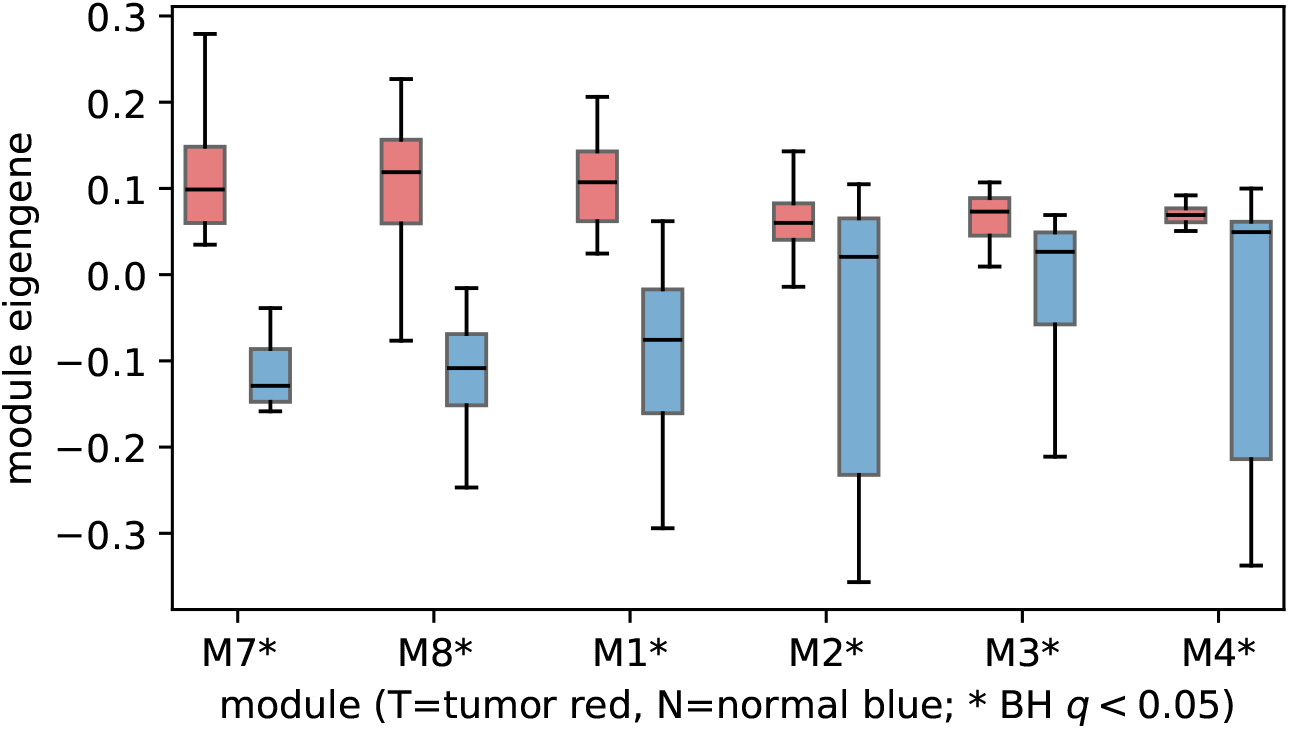
Module eigengenes separate tumour from normal tissue. Eigengene distributions (leading principal component of each module’s expression across the 60 samples) for the modules most associated with phenotype, split by tumour (T) and patient-matched normal (N) tissue; asterisks mark modules with a significant contrast (Benjamini–Hochberg *q <* 0.05).

### Visualizing cluster structure in reduced dimensions

To provide visual confirmation that the recovered modules occupy distinct regions of the space in which the objective operates, we embed each clustering result into two dimensions using PCA and correlation MDS (1 − |**R**|; see Methods). On Synthetic-Easy the three recovered modules form well-separated clouds in both embeddings (Fig 8), consistent with the perfect ARI. On Synthetic-Hard, where noise and unequal module sizes degrade separation, PCA shows considerable overlap because co-expressed but anti-correlated genes are geometrically distant; the correlation MDS, which measures distance in the same absolute-correlation space as the objective, reveals clearer module boundaries. For the empirical breast-cancer data, the correlation-MDS embedding of the 12 modules shows that even on real gene expression data the recovered modules occupy coherent, distinguishable regions of correlation space (Fig 9). These visualizations are produced automatically by the software and are available to users for any dataset.

**Fig 8.**
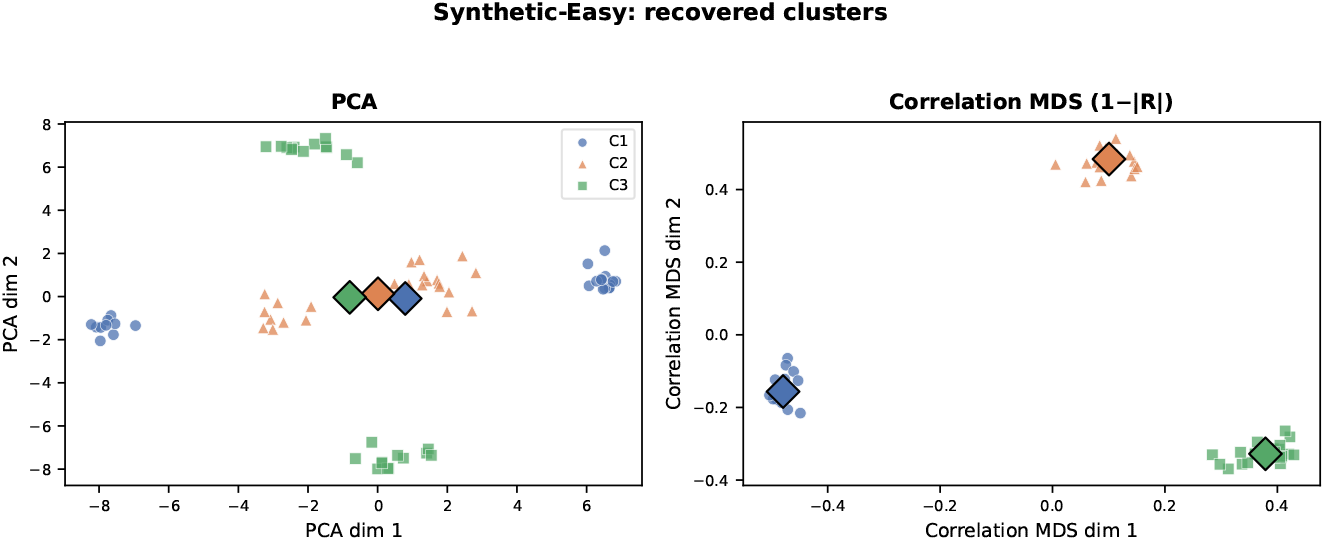
Dimensionality-reduction visualization of synthetic clustering. Recovered modules projected into two dimensions by PCA (left) and correlation MDS (right) on Synthetic-Easy. Diamonds mark cluster centroids. The correlation-MDS embedding uses the distance space native to the correlation objective and shows clean separation.

**Fig 9.**
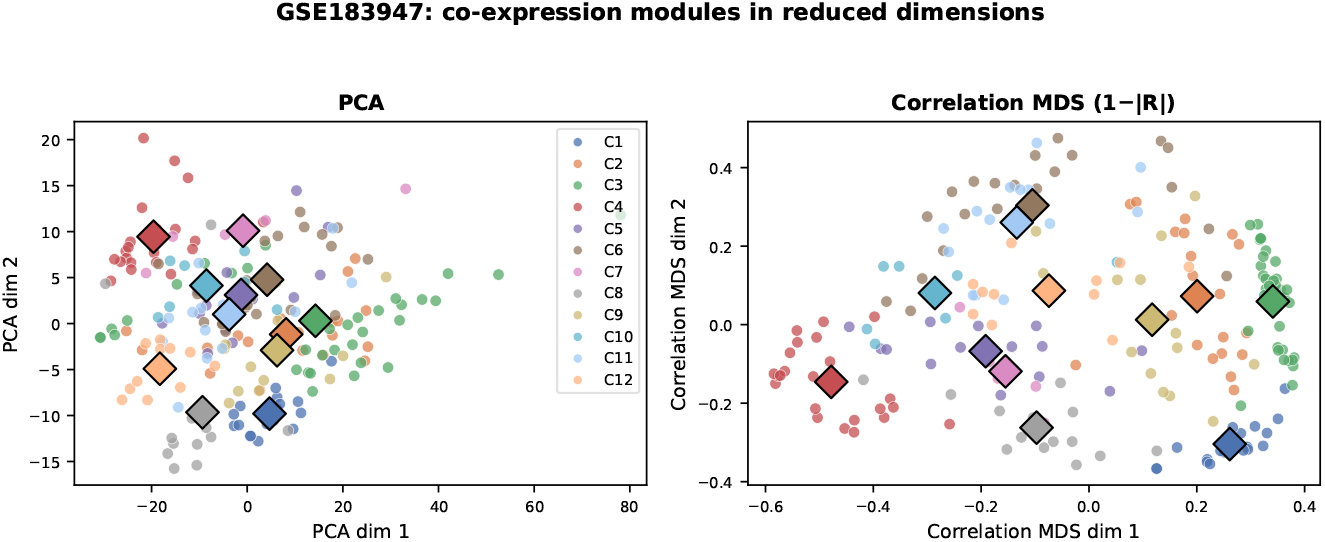
Dimensionality-reduction visualization of empirical modules. The 12 co-expression modules recovered from GSE183947 projected into two dimensions by PCA (left) and correlation MDS (right). Module boundaries are more distinct in the correlation distance space, confirming that the objective captures structure beyond what Euclidean geometry reveals.

## Discussion

GCM reframes co-expression module detection as maximum-likelihood inference under an explicit generative model. Solving that inference problem is globally effective. The correlation objective is not a heuristic: it is, up to a constant and the sample-size factor, the profile log-likelihood of a block-diagonal one-factor Gaussian model in which each module is driven by a single regulator with equal-magnitude *±* loadings [41] (S1 Appendix). This footing makes its model selection principled, in the sense that the number of modules is chosen by BIC/AIC over the *d* samples (Eq (3)), and explains why pairing it with a *geometric* information criterion fails: those assume Euclidean-Gaussian clusters, which the absolute-correlation model contradicts whenever modules contain anti-correlated genes.

Given the model, the contribution that matters in practice is optimization quality. Greedy agglomerative clustering on the correlation distance—the core of widely used co-expression pipelines (is a strong seed but a poor optimizer): once it merges two genes it can never separate them. Our replicated benchmark shows that hybridizing the GA with a memetic local search, which reassigns genes after the fact, surpasses hierarchical clustering across the noise range and does so consistently (lowest variance), and an ablation attributes most of the gain to the local-search step specifically. The same advantage holds when the number of modules is unknown and when unstructured genes must be ignored. In other words, the true (simulated) module structure is the likelihood optimum, and the value of the method is that it actually reaches it.

The approach has clear limits. Local search is what makes the optimizer competitive, so on problems where the objective does not decompose into incremental moves that are numerically inexpensive, it will be less efficient. Scaling to genome-wide matrices is bounded by the *O*(*n*^2^) correlation matrix and per-evaluation cost, so gene pre-filtering (as used here) is currently required. Natural extensions include richer within-module models (multi-factor or shrinkage-regularized free correlation, with the correspondingly larger parameter counts in the information criteria), correlation-aware crossover, and approximate or sketched correlation for very large *n*.

Methodologically, GCM sits in the tradition of GA-based and multi-objective clustering [19, 20] but is distinguished by a co-expression objective with an explicit likelihood, principled in-model selection, and a memetic optimizer that demonstrably is better than the greedy clustering such pipelines rely on—behind a single swappable-metric interface with a minimal dependency footprint. As a practical complement, the built-in dimensionality-reduction tools (PCA and correlation MDS, with optional t-SNE and UMAP) let users visually inspect and communicate the clustering structure, and the correlation-MDS embedding is particularly informative because it projects elements into the same distance space the objective operates in.

## Conclusion

We introduced GCM, a dependency-light Python tool that detects co-expression modules by maximizing the likelihood of an explicit block-diagonal one-factor Gaussian model with a memetic genetic algorithm. Placing the correlation objective on a likelihood footing makes its model selection principled, and hybridising the search with a greedy local refinement lets it reproducibly surpass the greedy agglomerative clustering on which standard co-expression pipelines rely. This advantage we establish with a replicated benchmark and an ablation, and that persists under unknown module count and unstructured-gene contamination. The same engine generalizes to geometric indices and recovers phenotype-associated modules in breast-cancer RNA-seq. The software, documentation, and full reproduction scripts are openly available.

## Supporting information

### S1 Appendix. Derivation of the correlation objective from the one-factor block model

#### Model

Each of the *d* samples (columns of **X**) is an independent draw **x** ~ *N* (**0**, Σ) over the *n* row-standardized genes, with Σ block-diagonal by the partition. Within a module *s* of size *n*_*s*_, the equal-magnitude one-factor structure *x*_*g*_ = *σ*_*g*_*f*_*s*_ + *ε*_*g*_, *σ*_*g*_ *∈ {±λ*_*s*_*}*, induces a correlation matrix that, after absorbing the loading signs by a diagonal *±*1 similarity (which changes neither eigenvalues nor determinant), is the equicorrelation (compound-symmetry) matrix Σ_CS_(*ρ*_*s*_) = (1 − *ρ*_*s*_)*I* + *ρ*_*s*_ **11**^*⊤*^ with 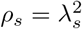. Its eigenvalues are 1 + (*n*_*s*_ − 1)*ρ*_*s*_ once and 1 − *ρ*_*s*_ with multiplicity *n*_*s*_ − 1, so log det Σ_CS_(*ρ*_*s*_) = log[1 + (*n*_*s*_ − 1)*ρ*_*s*_] + (*n*_*s*_ − 1) log(1 − *ρ*_*s*_).

#### Profile likelihood

For *d* i.i.d. samples with sample correlation **S**_*s*_ the per-module Gaussian log-likelihood is 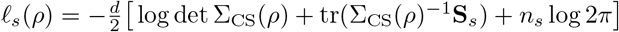. Writing *P* = **u**^*⊤*^**S**_*s*_**u** for the variance along the aligned direction 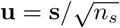 (sign vector **s**), the two distinct eigenvalues of Σ_CS_ are matched at their ML values 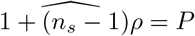, and 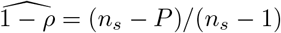 giving 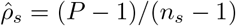. With sign alignment 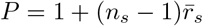 where 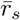 is the mean off-diagonal |*R*|, hence 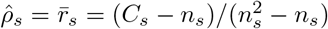 (Eq (1)). At the ML estimate the trace term collapses, tr 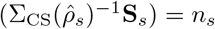, so 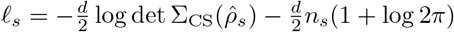.

#### Objective

Summing over modules, the second term is 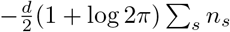, a constant (total genes fixed). The partition-dependent part is 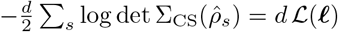, which is exactly *d* times Eq (2) after substituting 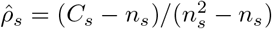 into the eigenvalue expression. The project’s objective is therefore the maximum log-likelihood of the model up to an additive constant and the factor *d*, and the per-module summand diverges to + *∞* as 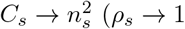, a perfectly correlated module); we cap this at a large finite value to keep the optimizer well-behaved.

#### Model selection

The *K*-module model has *K* free coherence parameters over *d* samples, giving BIC = − 2*d L* + *K* log *d* and AIC = − 2*d L* + 2*K* (Eq (3)). Because the likelihood carries the factor *d* and the penalty does not, the criteria license more modules as the number of samples grows, as required.

## Supporting information

**S1 File. Reproduction package**.For convenience, a compressed version of the repository is shared as a companion to the paper (although it is also available through GitHub and Zenodo). It contains the source code, the benchmark driver (`experiments/run_experiments.py`), and the generated results tables and figures.

## Data and Software Availability

All source code for the GCM package, along with the reproduction scripts to generate the results and figures presented in this manuscript, is publicly available under an open-source license on GitHub at https://github.com/luismadrigal98/GCM-MRK. To ensure long-term reproducibility, a persistent snapshot of the codebase at the time of submission has been archived on Zenodo (DOI: 10.5281/zenodo.21361265). The empirical breast cancer RNA-seq dataset (GSE183947) analyzed in this study is publicly available from the NCBI Gene Expression Omnibus (GEO).

## Acknowledgments

We thank all the class of Simulation in Python offered in the University of Kansas, where all this project of evolving solutions using evolutionary algorithms (the simulation bit) started.

